# Structure of the space of taboo-free sequences

**DOI:** 10.1101/824847

**Authors:** Cassius Manuel, Arndt von Haeseler

## Abstract

Models of sequence evolution typically assume that all sequences are possible. However, restriction enzymes that cut DNA at specific recognition sites provide an example where carrying a recognition sequence can be lethal. Motivated by this observation, we studied the set of strings over a finite alphabet with **taboos**, that is, with prohibited substrings. The taboo-set is referred to as 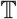 and any allowed string as a taboo-free string. We consider the graph 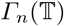 whose vertices are taboo-free strings of length *n* and whose edges connect two taboo-free strings if their Hamming distance equals 1. Any (random) walk on this graph describes the evolution of a DNA sequence that avoids deleterious taboos. We describe the construction of the vertex set of 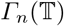. Then we state conditions under which 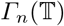 and its suffix subgraphs are connected. Moreover, we provide a simple algorithm that can determine, for an arbitrary 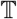, if all these graphs are connected. We concluded that bacterial taboo-free Hamming graphs are nearly always connected, although 4 properly chosen taboos are enough to disconnect one of its suffix subgraphs.

## 1 Introduction

In bacteria, restriction enzymes cleave foreign DNA to stop its propagation. To do so, a double-stranded cut is induced by a so-called recognition sequence, a DNA sequence of length 4 – 8 base pairs ([2]). As part of their recognition-modification system, bacteria can escape the lethal effect of their own restriction enzymes by modifying recognition sequences in their own DNA ([14]). Nevertheless, a significant avoidance of recognition sequences has been observed in bacterial DNA ([7], [19]). Also in bacteriophages, the avoidance of the recognition sequences is evolutionary advantageous ([19]). Therefore the recognition sequence is, as we call it, a **taboo** in a majority of fragments of both host and foreign DNA.

Motivated by this biological phenomenon, we studied the Hamming graph 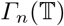, whose vertices are strings of length *n* over a finite alphabet *Σ* not containing any taboo of the set 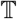 as substring. By definition, an edge connects two strings of this graph if their Hamming distance is 1, i.e. if they differ by a single substitution. Moreover, given a taboo-free string *s*, we consider the subgraph 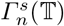 of 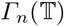 induced by every taboo-free string with suffix *s*. Suffix *s* represents a conserved DNA fragment, that is, a sequence which remained invariable during evolution ([20], [6]).

Instead of studying just the connectivity of 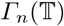, here we define conditions under which every graph of the form 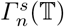 is connected. Note that **this result is stronger than just the connectivity of** 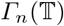, for 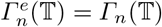, where *e* is the empty string. If taboo-set 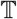 satisfies that, for every taboo-free string *s* and integer *n*, graph 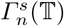 is connected, then evolution can explore all the space of taboo-free sequences by simple point mutation, no matter which DNA suffix fragments remain invariable. Notice that we assume that the taboo-set 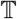 does not change in the course of evolution.

## 2 Spaces of taboo-free sequences

We start with a non-biological example of a space of taboo-free strings. For the binary alphabet *Σ* = {0, 1} and 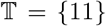, graphs 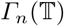 are known as Fibonacci cubes, proposed in [9] and [10] as a network topology in parallel computing. Fibonacci cubes are connected, bipartite and contain a Hamiltonian path. Enumeration results regarding e.g. the Wiener Index or the degree distribution have been obtained, mainly using generating functions, in [13], [5] and [16], among others (see the compendium [12]).

Other types of taboo strings assuming a binary alphabet *Σ* = {0, 1} have been considered in [10] for 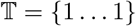, while the case 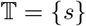 for an arbitrary binary string *s* was studied in [11]. The number of strings over any finite alphabet with taboos has been studied, using generating functions, in [8], while related software is given in [17].

Therefore our terminology will be based on this previous work: From now on we will use the terms **string** to refer to a sequence of symbols over an alphabet *Σ*, while **(DNA) sequence** is reserved for biological contexts, where the alphabet consists of the four nucleotides, i.e. *Σ* = {*A, C, G, T*}. Given a string *s*, its length is denoted by |*s*|. Given two strings *s*_1_, *s*_2_ of equal length, then *d*(*s*_1_, *s*_2_) denotes their Hamming distance, that is, the number of sites at which the corresponding symbols differ.

Given a finite set of strings 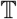, we call 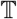 a **taboo-set** if, for any 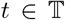, |*t*| ≥ 2. We refer to the strings of 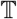 as **taboos**. The length of the longest taboo in 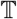 will be denoted by 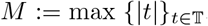. A **taboo-free string** is a string not having any taboo of 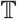 as substring. The set of taboo-free strings of length *n* is denoted by 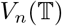. Given a taboo-free string *s*, for 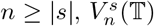 denotes the set of strings of 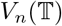 with suffix *s*.

**The Hamming graph of taboo-free strings of length** *n*, denoted by 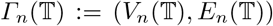, is the graph with vertex set 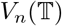 such that two vertices 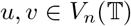 are adjacent if their Hamming distance equals 1. See Fig. 1 for an example where 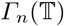 is disconnected for *n* ≥ 3. Analogously, 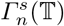 is the Hamming graph with vertex set 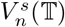.

**Fig. 1.**
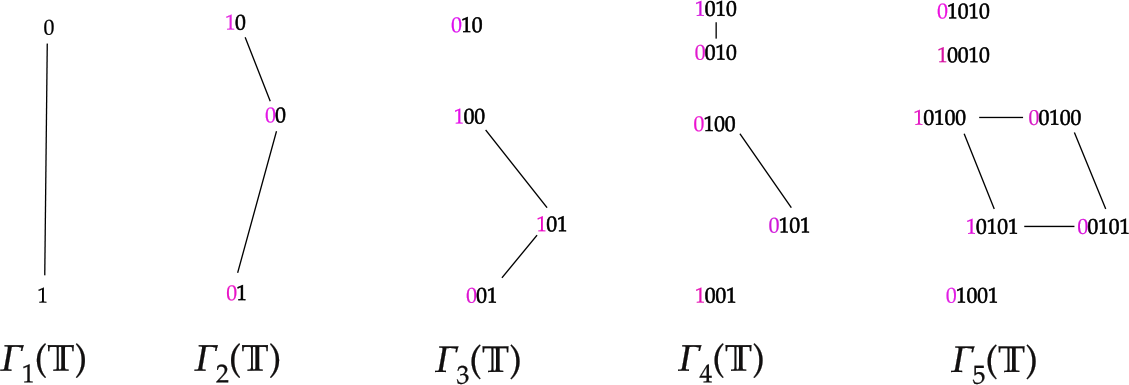
Graph 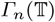 for *n* ∈ [1, 5] for binary alphabet *Σ* = {0, 1} and 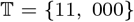. Set 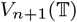 is constructed by adding every allowed symbol at the beginning of each string in 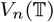.

Taboo-sets as generated by the avoidance of restriction sequences can assume various levels of complexities. We discuss some examples from REBASE ([18]). Note that many restriction enzymes of this database have an unknown recognition sequence, hence our taboo-sets may underestimate the actual amount of taboos. Before studying the examples, we will briefly review essential nomenclature for DNA sequences.

DNA is double-stranded, where *A* pairs with *T* and *G* pairs with *C*, thus it suffices to discuss only one of the strands. We adopt the convention that, given any of the strands, the DNA sequence is always represented from the 5’ end to the 3’ end (which is chemically determined). As a consequence, given a DNA sequence, **its complementary DNA sequence**, the one lying on the opposite strand, is obtained by inverting the order of the symbols and carrying through substitutions *A ↔ T* and *C ↔ G*. If a DNA sequence *s* is identical to its complementary DNA sequence, we say that *s* is an **inverted repeat** (see [22]). For example, sequence *CCGG* is an inverted repeat.

The fact that DNA is double-stranded implies that each recognition sequence induces taboos in pairs, namely itself and its complementary DNA sequence. For example, if *AGGGC* is a recognition sequence, then also the complementary strand *GCCCT* is a taboo. If, however, the recognition sequence is an inverted repeat such as *TGCA*, then this pair is actually one single recognition sequence. This is the case in most of type II restriction-modification systems ([7]).

### 2.1 Turneriella parva

The *Turneriella parva* (REBASE organism number 8970) strain produces a restriction enzyme with recognition sequence *GATC*, an inverted repeat. Similarly, another of its enzymes has recognition sequences *GGACC* and *GGTCC*. Thus, these restriction enzymes generate the taboo-set

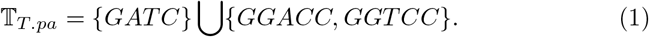

### 2.2 Helicobacter pylori

In *H. pylori* 21-A-EK1 (studied in [1]), many restriction enzymes have been identified. For the sake of clarity, let us write 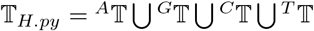, where 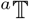 denotes those taboos in 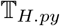 whose **first** symbol is *a* ∈ *Σ*. Then we have

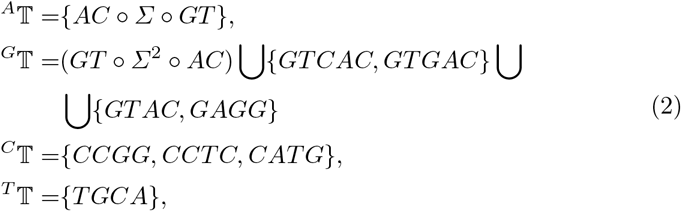

where *GT ○ Σ*^2^ ○ *AC* represents taboos of the type *GGabCC* with *a, b* ∈ *Σ*, and so on for analogous notations. In Section 10 we will show that every graph 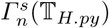 is connected. Thus, in particular, 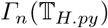 is connected.

### 2.3 An imaginary bacterium

The taboo-set can significantly influence evolution in the cases where some 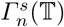 is disconnected. To explain this, we will create a plausible, nonexistent example. Suppose that a strain of *Bacterium imaginara* has taboo-set

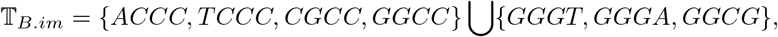

where the second set contains the complementary DNA sequences of the first set, except that of *GGCC*, which is an inverted repeat. At first glance, tabooset 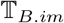 seems less restrictive than 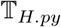, which has 6 taboos of length four and 22 taboos of length five or more. However, we will show in Section 10 that the graph 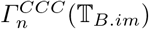 is disconnected for *n* ≥ 5. This happens because the first set in 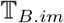 represents a “wall of taboos” between graphs 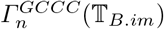 and 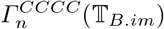.

This produces the following evolutionary implications: Assume that we have two correctly aligned DNA fragments *f_α_* and *f_β_* from viruses *α* and *β* that infect *Bacterium imaginara*. Assume moreover that we can write *f_α_* = *r_α_GCCC* and *f_β_* = *r_β_CCCC* for some strings *r_α_* and *r_β_*, as also that the suffix *CCC* is invariable due to functional constrains. Then *f_α_* cannot have evolved from *f_β_* by simple point mutations, because at some point in evolution a taboo string is produced, what would be lethal for the carrier. Thus, the standard models of sequence evolution do not apply ([21]).

Thus, taboo-set 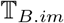 is an interesting source of information to study evolution. This motivates our study, from a general point of view, of the Hamming graph with taboos.

## 3 Outline of the theory of 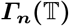

We will characterize those taboo-sets 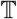 such that every graph of the form 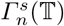 is connected. To do this we introduce a very general type of taboo-sets, which we call **left proper** (Sec. 5, Def. 4). Left proper taboo-sets satisfy that, for each string 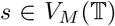, a symbol *a* ∈ *Σ* exists such that *as* is taboo-free. Our biological examples discussed so far are left proper.

For the sake of clarity, consider the DNA-alphabet *Σ* = {*A, C, G, T*} and the taboo-set 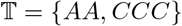. Since 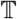 is left proper, for any taboo-free string *s* of length at least *M* = 3 there exists a string *w* of any desired length such that *ws* is taboo-free (Def. 3 of k-prefix, Prop. 5 and Prop. 6).

Then **synchronization** comes into play (Def. 6). Two taboo-free strings *s*_1_ and *s*_2_ are left *k*-synchronized if there exists a string *w* of length *k* such that *ws*_1_ and *ws*_2_ are taboo-free. More informally, synchronization sticks together connected components that partition the Hamming graph. Prop. 7 states that, if *s*_1_ and *s*_2_ are left (*M* − 1)-synchronized, then they are left *k*-synchronized for every *k* ∈ ℕ. For taboo-set 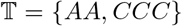, given two taboo-free strings *s*_1_ and *s*_2_, it holds that *TTs*_1_ and *TTs*_2_ are taboo-free. Thus *s*_1_ and *s*_2_ are left *k*-synchronized for every *k* ∈ ℕ.

In Sec. 6 and 7, we realize that not the whole string *s* is relevant to construct graph 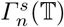, but only the biggest prefix of *s* which is a suffix of a taboo, which we call *s*[1, *k_s_*] (Prop. 10). In particular, we infer the graph isomorphism 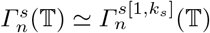 (Theorem 15). This implies that, to study every graph 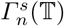, it is enough to consider 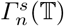 for *s* ∈ {*e, A, C, CC*}, where *e* is the empty string.

In Section 8 we introduce **the quotient graph**, a useful tool to describe the structure of graphs with many vertices. Given a partition of the vertices, the quotient graph represents the edges connecting the sets of this partition. Lemma 19 states that, if 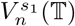 and 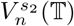 are two of the sets of this partition, then they are adjacent in the quotient graph for any *n* ∈ ℕ iff *d*(*s*_1_, *s*_2_) = 1 and *s*_1_ and *s*_2_ are left (*M* − 1)-synchronized. For 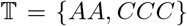, every 2 strings are left 2-synchronized, implying that only condition *d*(*s*_1_, *s*_2_) = 1 is needed.

Combining all these results, we can prove the connectivity of any 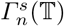 (Sec. 9). This is done as follows: We check first if 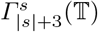 is connected for any *s* ∈ {*e, A, C, CC*}. Using the classification of Theorem 15, that implies that, for any 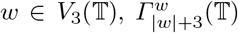 is connected. Then we repeatedly make use of the quotient graph induced by partition 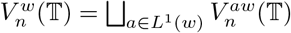 for 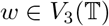, which allows us to inductively prove that every 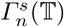 is connected (Lemma 20 and Theorem 21).

Remarkably, connectivity of the biological examples we have dealt with (as also of 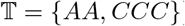) can be proven by applying Prop. 24, which does not make use of most of the theory we developed. In fact, only the definition of *k*-suffix and left *k*-synchronization are sufficient to understand and apply Prop. 24. However, our theory can handle a much more general types of taboo-sets.

## 4 Basic notation and properties of set 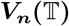

Our alphabet comprises *m* ≥ 2 symbols *Σ* = {*a*_1_, · · ·, *a_m_*}. The set of strings over *Σ* of length *n* will be referred to as *Σ^n^*, with cardinality |*Σ^n^*| = *m^n^*. The empty string will be denoted by *e*, and satisfies |*e*| = 0 and {*e*} = *Σ*^0^.

If string *s*_1_ is substring of string *s*_2_, we write *s*_1_ ≺ *s*_2_, while *s*_1_ ⊀ *s*_2_ denotes that *s*_1_ is **not** a substring of *s*_2_. By convention, *e* ≺ *s* for any string *s*. Given two strings *s*_1_ and *s*_2_, we define *s*_1_*s*_2_ as the concatenation of *s*_1_ and *s*_2_. Note that *es* = *se* = *s* for any *s*. Given a set of strings *S* = {*s*_1_, · · · *s_k_*} and a string *s*, we define *s* ○ *S* := {*ss*_1_, · · · *ss_k_*} and *S* ○ *s* := {*s*_1_*s*, · · · *s_k_s*}. Given two string sets *S*_1_ and *S*_2_, we define *S*_1_ ◦ *S*_2_ := {*s*_1_*s*_2_}_*s*_1_ ∈ *S*_1_; *s*_2_ ∈ *S*_2__, and 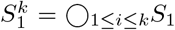. If *S*_1_ and *S*_2_ are disjoint, then the disjoint union of *S*_1_ and *S*_2_ will be denoted by 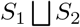.

### Definition 1

*Given symbols b*_1_, · · ·, *b_n_* ∈ *Σ, let us consider string s* = *b*_1_ · · · *b_n_. For i* ∈ ℕ *and j* ∈ ℕ_0_, *we define*

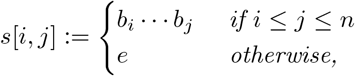

*i.e. s*[*i, j*] *denotes the substring of s starting at the i-th symbol up to the j-th symbol, and e when this substring is not well-defined (for example if j* = 0*). Given a set of strings S, we define*

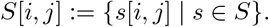

### Example 1

*If Σ* = {*D, O, R, W*} *and s* = *WORD, then we have s*[1, 1] = *W, s*[1, 2] = *WO, s*[2, 4] = *ORD and s*[1, 0] = *e*.

### Proposition 1

*For any two taboo-sets* 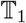 *and* 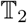 *and any n* ∈ ℕ, *it holds that*:

a. *Set* 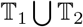 *is a taboo-set and* 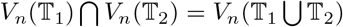.
b. *If for every* 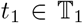 *there exists* 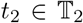 *such that t*_2_ ≺ *t*_1_, *then for any* 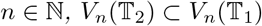.

*Proof*

a. Every 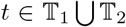 has length at least 2, thus 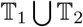 is a taboo-set. The fact that a string 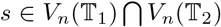 satisfies *t*_1_ ⊀ *s* for any 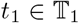 and *t*_2_ ⊀ *s* for any 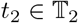 is equivalent to *s* satisfying *t* ⊀ *s* for any 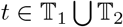.
b. Consider 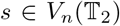. Assume that 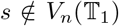 then there exists 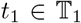 such that *t*_1_ ⊀ *s*. But there also exists a 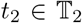 such that *t*_2_ ≺ *t*_1_, thus *t*_2_ ≺ *s*, a contradiction. Hence 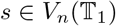.

The set 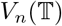 can be obtained from other taboo-sets than 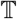. In this sense, taboo-sets are not unique, as we illustrate in the following proposition.

### Proposition 2

*Given string s, for any n* ≥ 1 + |*s*| *it holds that*

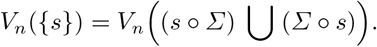

*Proof*

−⊆: Any taboo in 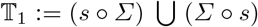 has 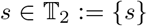 as substring, thus Prop. 1.*b* implies 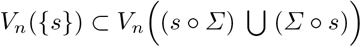.
−⊇: Assume there is 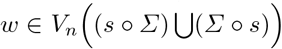 with *s* ≺ *w*. Since |*w*| = *n* and *n* ≥ |*s*| + 1, the substring *s* is either preceded or followed by some symbol *a* ∈ *Σ*. This contradicts {*as, sa*} ⊂ (*s* ○ *Σ*) ∪ (*Σ* ○ *s*).

Prop. 2 implies that, for any 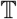, we can construct many taboo-sets 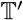 such that 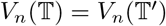 for any *n* ≥ max(*M, M′*), where *M′* denotes the length of the longest taboo in 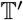.

### Example 2

*If* 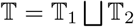 *with* 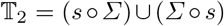, *Prop. 1.a and 2 imply that* 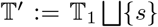 *satisfies* 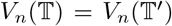 *for any n* ≥ *M. Repeating this process, we can construct a taboo-set* 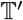 *such that* 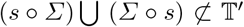 *for any string s and satisfying* 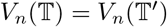 *for any n* ≥ *M*.

Example 2 motivates the following definition.

### Definition 2

*A taboo-set* 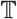 *is **minimal** if the following conditions hold*:

– *For every j* ∈ [0, *M* − 1] *and* 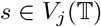, *we have* 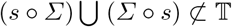.
– *For every different* 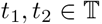, *it holds that t*_1_ ⊀ *t*_2_.

When a taboo-set 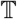 is not minimal, it can be understood that this 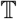 is unnecessarily complicated. For computational purposes, it is recommendable to force 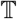 to be minimal. Apart from this, also *M*, the length of the longest taboo, is of great importance: If *M* is known, the following result guarantees that the concatenation of three taboo-free strings is a taboo-free string.

### Proposition 3

*Given* 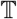, *consider strings s*_1_, *s*_2_, *and s*_3_ *of respective lengths j*_1_, *j*_2_, *j*_3_ ∈ ℕ_0_. *If* 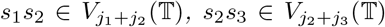 *and j*_2_ ≥ *M* − 1, *then* 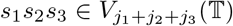.

*Proof* Assume either *j*_1_ > 0 or *j*_3_ > 0, yielding *j* := *j*_1_ + *j*_2_ + *j*_3_ ≥ *M*. For each *i* ∈ [1, *j* + 1 − *M*], the fact that *j*_2_ ≥ *M* − 1 implies that either *s*[*i, i* + *M* − 1] ≺ *s*_1_*s*_2_ or *s*[*i, i* + *M* − 1] ≺ *s*_2_*s*_3_, hence each *s*[*i, i* + *M* − 1] is taboo-free and the result follows.

## 5 Prefixes and suffixes of a taboo-free string

Given a taboo-free string *s*, the construction of set 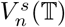 for *n* > |*s*| depends on which strings can be concatenated to the left side of *s*, yielding 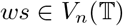. This object is of interest for us.

### Definition 3

*For taboo-set* 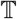, *j* ∈ ℕ_0_ *and string* 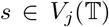, *let k* ∈ ℕ_0_ *be given. The k**-prefixes** of s are the elements of the set L^k^*(*s*), *defined as*

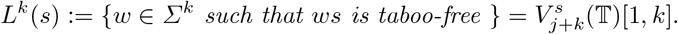

*If L^k^*(*s*) ≠ ∅, *then we will say that s is k**-prefixable**. Similarly, the k**-suffixes** of s, denoted R^k^*(*s*), *are the strings w* ∈ *Σ^k^ such that* 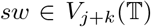. *When R^k^*(*s*) ≠ ∅, *we say that s is k**-suffixable***.

### Example 3

*If Σ* = {0, 1} *and* 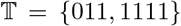, *then L*^1^(11) = {1}, *while L*^2^(11) = ∅*. Hence string* 11 *is* 1*-prefixable, but not* 2*-prefixable. Moreover, R*^1^(11) = {0, 1}, *hence string* 11 *is* 1*-suffixable*.

By construction, given 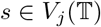, for any *n* ∈ ℕ_0_ it holds that

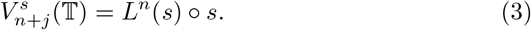

That is, 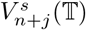 is *L^n^*(*s*) with suffix *s* appended. Moreover, the *k*-prefixes of a string *s* induce a partition of the set 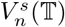.

### Proposition 4

*Given a taboo-set* 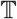, *an integer k* ∈ ℕ_0_ *and a string* 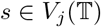 *with j* ∈ ℕ_0_, *for any n* ≥ *j* + *k it holds that*

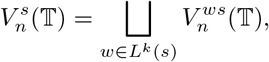

*that is, the set* 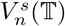 *can be partitioned into the disjoint sets of taboo-free strings of length n with suffix ws, where w* ∈ *L*^*k*^(*s*).

*Proof* If *s* is not *k*-prefixable, then *L^k^*(*s*) = ∅ and 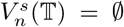, hence the equation holds. Otherwise, the inclusion ⊃ is clear, while the ⊂ follows from the fact that, for any string *w* ∈ *Σ^k^* preceding the suffix *s*, this *w* must necessarily belong to *L^k^*(*s*).

Clearly, if a taboo-free string 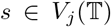 is *k*^∗^-prefixable, then it is also *k*-prefixable for any integer *k < k*^∗^, while nothing can be said *a priori* about the case *k > k*^∗^. We can, however, define a type of taboo-sets satisfying that, if *s* has length at least *M*, then *s* is *k*-prefixable for any *k* ∈ ℕ.

### Definition 4

*A taboo-set* 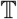 *is **left proper** if every* 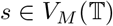 *is* 1*-prefixable. Analogously*, 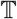 *is **right proper** if every* 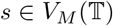 *is* 1*-suffixable.*

### Example 4

*If Σ* = {*A, C, G, T*} *and* 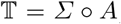, then 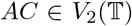 *and AC is not* 1*-suffixable. Thus* 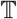 *is not left proper*.

### Proposition 5

*Consider a left proper taboo-set* 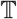, *j* ∈ ℕ *and* 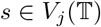, *such that either of the following conditions holds*.

a. *j* ≥ *M*
b. *j* ≤ *M* − 1 *and s is* (*M* − *j*)*-prefixable*

*Then s is k-prefixable for any k* ∈ ℕ.

*Proof* If condition *a*) applies, then consider 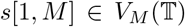, yielding that *s* is 1-prefixable, that is, there exists *a* ∈ *Σ* with 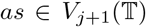. Proceeding analogously with (*as*)[1, *M*], we would infer that *s* is 2-prefixable. Continuing with this process, we deduce that *s* is *k*-prefixable for any *k* ∈ ℕ.

If condition *b*) holds, then we can take any string in 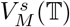 and proceed as we did assuming *a*).

Prop. 5 is the reason why we only study proper taboo-sets, for the existence of arbitrary *k*-suffixes is necessary in many of our proofs. In particular, we will mostly focus on the case in which 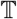 is left proper, although analogous results for right proper taboo-sets are obtained by reversing the order of the symbols composing the string.

When the length of a taboo-free string is at least *M*, then Prop. 5 is easy to apply. Otherwise, one needs to study the *k*-prefixability of this string. To that end, the following definition can be useful.

### Definition 5

*For set of strings S, **the suffixes of** S are defined as*

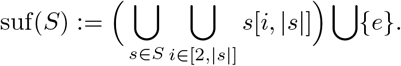

### Example 5

*If S* = {*ACG, GGG, TTC, CC*} *then*

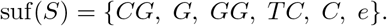

If 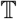 is left proper, then 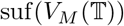 is of great interest, for it indicates which small suffixes are prefixable, as the following proposition indicates.

### Proposition 6

*Consider a left proper taboo-set* 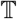, *j* ∈ ℕ_0_ *and* 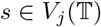.

a. *If j* ≤ *M* − 1 *and* 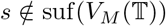, *then for every* 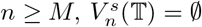.
b. *If either j* ≥ *M or* 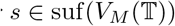, *then for n* ≥ max(*j, M*), 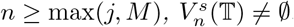.

*Proof*

a. If *j* ≤ *M* − 1 and 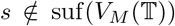, since 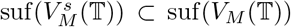, it holds that 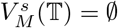. This implies that 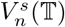 is empty for every *n* ≥ *M*, because otherwise

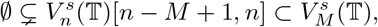

what would be a contradiction.
b. If *j* ≥ *M*, since 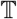 is left proper, Prop. 5 does the work. If 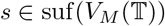, then *s* is (*M* − *j*)-prefixable, so again we apply Prop. 5.

To prove the connectivity of graphs 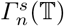, we will need to know whether two different strings have a *k*-prefix in common, hence a definition is needed.

### Definition 6

*Given a taboo-set* 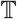, *we say that two taboo-free rings s and r (maybe of different length) are **left** k**-synchronized** if* 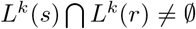. *If* 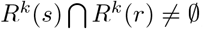, *then we say that s and r are **right** k**-synchronized***.

Clearly, two taboo-free strings that are left *k*^∗^-synchronized are also left *k*-synchronized for any *k* ≤ *k*^∗^ (one simply has to “cut” the last *k* symbols of 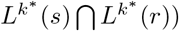. The following proposition states when we can also guarantee this property for *k > k*^∗^ if 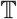 is left proper.

### Proposition 7

*Consider a left proper taboo-set* 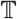 *and two taboo-free strings s, r, with length greater than zero, such that s and r are left* (*M* −1)*-synchronised. Then s and r are left k-synchronized for any k* ∈ ℕ.

*Proof* Clearly the case *k* ≤ *M* − 1 holds. For *k > M* − 1, consider a string *w* 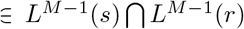. We know that *ws* and *wr* are taboo-free strings with length at least *M*. Since 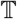 is left proper, Prop. 5 implies that *w* is *k′*-prefixable for any *k′* ∈ ℕ. For any *k′*, take *x* 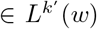, and consider strings *xws* and *xwr*. The fact that |*w*| = *M* − 1, together with the fact that *xw* and the pair *ws, wr* are taboo-free, allows applying Prop. 3, hence *xws* and *xwr* are also taboo-free. All in all, 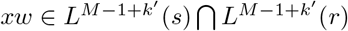. We write *k* := *M* − 1 + *k′*, yielding the result for any *k > M* − 1.

For cases where the number of taboos is small, the following proposition is an easy way to prove that every two taboo-free strings of length *M* are left *k*-synchronized.

### Proposition 8

*Take a left proper taboo-set* 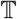 *such that any* 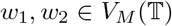 *with d*(*w*_1_, *w*_2_) = 1 *are left* 1*-synchronized. Then, for each* 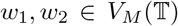 *with d*(*w*_1_, *w*_2_) = 1, *w*_1_ *and w*_2_ *are left k-synchronized for any k* ∈ ℕ_0_.

*Proof* Given any left 1-synchronized pair *w*_1_, *w*_2_ with *d*(*w*_1_, *w*_2_) = 1, there exists *a* ∈ *Σ* such that *aw*_1_ and *aw*_2_ are taboo-free. Since 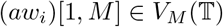 for *i* ∈ {1, 2} and the Hamming distance between these two strings is at most 1, *aw*_1_, *aw*_2_ are 1-synchronized, hence there exists *b* ∈ *Σ* such that *baw*_1_ and *baw*_2_ are taboo-free, i.e. *w*_1_ and *w*_2_ are left 2-synchronized. Continuing with this process, the result follows.

## 6 Simplification of suffixes

Consider two taboo-free strings *w*, *s* and the set 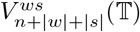. In general, it is not true that 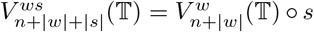, because it can happen that, for some 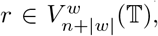, the concatenation *rs* creates a taboo substring. Only if 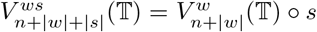, one can simply take 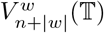 an append an *s* at the end of each of its strings. This property is studied in this section.

### Proposition 9

*For a taboo-set* 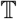, *consider w* ∈ *Σ^M^*^−1^ *and s* ∈ *Σ^j^, j* ∈ ℕ_0_, *such that* 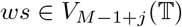. *Then for any n* ≥ *M* − 1,

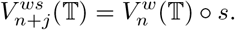

*Proof* The inclusion ⊂ is clear. The inclusion ⊃ follows from the fact that, if we are given 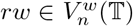 such that 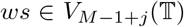, since |*w*| = *M* − 1, we can apply Prop. 3, yielding that *rws* is a taboo-free string.

### Definition 7

*Given a taboo-set* 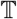 *and* 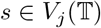, *j* ≥ 0, *we define*

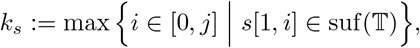

*i.e. k_s_ denotes the length of the longest prefix of s being the suffix of a taboo*.

Note that *k_s_* is always well defined, for 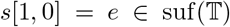, hence *k_s_* ∈ [0, min (*M* − 1, *j*)]. Using Def.7, Prop. 9 can be improved.

### Proposition 10

*Let a taboo-set* 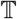, *j* ∈ ℕ_0_ *and* 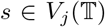 *be given. For any n* ∈ ℕ_0_ *it holds that*

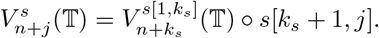

*Proof* The result is obvious if *j* = 0 or *n* = 0, hence assume *j* > 0 and *n* > 0. Along the reasoning that follows, note that it also applies for the case *k_s_* = 0. Clearly 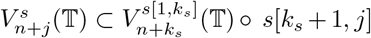. For 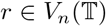, consider 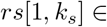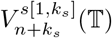. We need to prove that the string

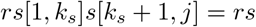

is taboo-free. But otherwise, since *rs*[1, *k_s_*] and *s* are taboo-free, there would exist integers *c, d* such that 1 ≤ *c ≤ |r| ≤ r* + *k_s_ < d ≤ |r|* +*j* and 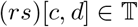. Take *k*^∗^ := *d − |r| > k_s_*, what yields 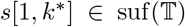, contradicting the maximality of *k_s_*. Hence *rs* is taboo-free, as desired.

### Corollary 11

*Given a taboo-set* 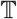, *j* ∈ ℕ_0_ *and* 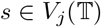, *then for any k* ∈ ℕ_0_

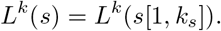

*Proof* By construction, 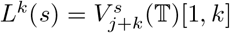. Prop. 10 yields

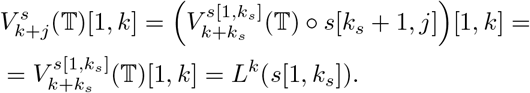

### Corollary 12

*Given taboo-set* 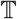 *and any pair of taboo-free strings s and r, the following statements are equivalent for any k* ∈ ℕ_0_:

– *s and r are left k-synchronized*
– *s*[1, *k_s_*] *and r*[1, *k_r_*] *are left k-synchronized*.

*Proof* Strings *s* and *r* are left *k*-synchronized iff 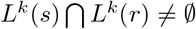. We just have to apply Corollary 11.

All in all, we learnt in this section that string *s*[1, *k_s_*] is informative enough to construct 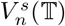 or *L^k^*(*s*).

## 7 Isomorphisms of graphs 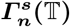

We will make use of common terminology of graph theory (cf. [24]). In this work, the term “graph” denotes a simple undirected graph. We say that graph *G*_1_ = (*V*_1_, *E*_1_) is **subgraph** of *G*_2_ = (*V*_2_, *E*_2_) if *V*_1_ ⊂ *V*_2_ and *E*_1_ ⊂ *E*_2_, and write *G*_1_ ⊂ *G*_2_.

Given a graph *G* = (*V, E*) and a subset *S* ⊂ *V*, the **subgraph induced by** *S* in *G*, *G*(*S*) = (*S*,*E*_*S*_), has vertex set *S* and, for any *u, v ∈ S*, {*u, v*} ∈ *E_S_* iff {*u, v} ∈ E*. To avoid a too complex notation, given a subset 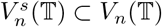 for some taboo-free string *s*, we refer to the subgraph induced by 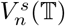 in 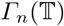 as 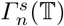. In particular, 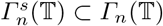.

Two graphs *G*_1_ = (*V*_1_, *E*_1_) and *G*_2_ = (*V*_2_, *E*_2_) are **isomorphic**, denoted by *G*_1_ ≃ *G*_2_, if there exists a bijection *f* : *V*_1_ → *V*_2_ such that, for every *u, v* ∈ *V*_1_, {*u, v*} ∈ *E*_1_ iff {*f* (*u*), *f* (*v*)} ∈ *E*_2_. That is, *G*_1_ and *G*_2_ are isomorphic if there exists an edge-preserving bijection between their vertex sets.

### Proposition 13

*Given* 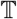, *j* ∈ ℕ_0_ *and* 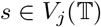, *consider a taboo-free string w satisfying* 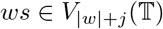 *and such that, for a given n* ≥ |*w*|,

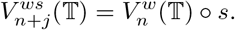

*Then* 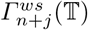 *and* 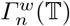 *are isomorphic*.

*Proof* Since the vertex set of 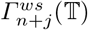 is 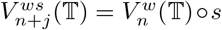, we will establish this bijection using the map

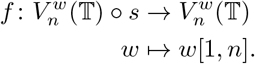

The map *f* is well defined and bijective. Moreover, this is an edge-preserving bijection: Given any pair of strings *s*_1_, *s*_2_ ∈ *Σ*^*n*^ and any string *s* ∈ *Σ*^*j*^, *d*(*s*_1_, *s*_2_) = 1 iff *d*(*s*_1_*s, s*_2_*s*) = 1.

Propositions 13 and 9 imply that, given 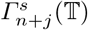 for some 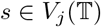 with *j* ≥ *M*, we can do 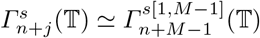. Going further, Prop. 10 yields

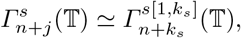

or differently said, we have split graphs 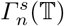 into classes, each of which is represented by 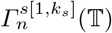:

### Proposition 14

*Consider a taboo-set* 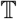, *j* ∈ ℕ_0_ *and* 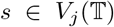. *Then a unique* 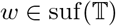 *exists such that w* = *s*[1, *k*_*s*_]. *Moreover, for any n* ≥ 0,

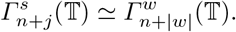

Prop. 14 does not describe in which cases 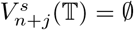. However, if 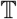 is left proper, Prop. 6 implies that this happens iff *j* ≤ *M* − 1 and 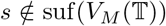. This suggests that we can state a version of Prop. 14 for left proper 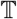, although before doing so let us introduce a new object.

### Definition 8

*Given a left proper taboo-set* 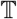, ***the long suffix classification*** 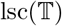 *is defined as*

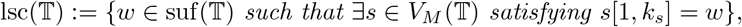

*that is*, 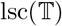 *is the set of all suffixes of taboos that are the longest prefix of at least one taboo-free string of length M*.

### Example 6

*If Σ*_1_ = {*A, C, G, T*} *and* 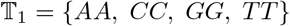, *then*

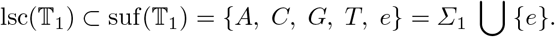

*For any* 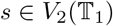, *we see k*_*s*_ > 0, *hence* 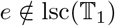. *Moreover*,

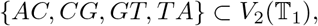

*yielding* 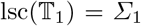. *If we consider Σ*_2_ := {*A, C, G, T, C′*}, *where C′ could represent a 5-methylcytosine, and* 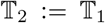, *then string s* = *C′ A satisfies k*_*s*_ = 0, *hence* 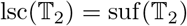.

The following theorem classfies graphs 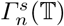 for left proper 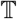.

### Theorem 15

*Given a left proper taboo-set* 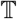 *and j* ∈ ℕ_0_, *consider* 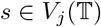 *such that either j* ≥ *M or* 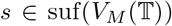. *Then* 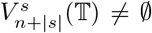 *for n* ≥ 0. *There exists a unique w* 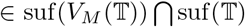 *such that w* = *s*[1, *k*_*s*_], *which satisfies* 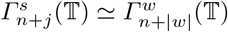 *for n* ≥ 0. *If j* ≥ *M, then* 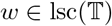.

*Proof* Prop. 6 yields 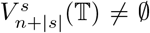 for *n* ≥ 0, and 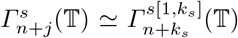 for *n* ≥ 0 follows from Prop. 14. Hence we can set *w* := *s*[1, *k*_*s*_], which by definition belongs to 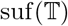. We also have 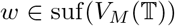, which follows from *s* being *k*-suffixable for any *k* ∈ ℕ_0_ and Prop. 6. This *w* is trivially unique since *k*_*s*_ is uniquely determined given *s*. If *j* ≥ *M*, the fact that 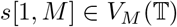 and the definition of 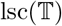 imply that 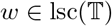.

To compute 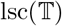, we recommend that 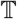 be minimal, for otherwise the complexity of the problem increases. Theorem 15 implies that

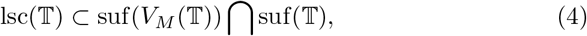

thus we define the **short suffix classification** as

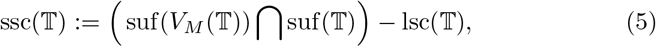

or equivalently,

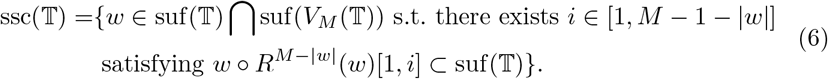

Eq. 6 is quite technical, but if *w* satisfies the stronger condition |*w*| < *M* − 1 and 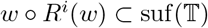 for some *i* ∈ [1, *M* − 1 − |*w*|], then any *s* ∈ *w* ○ *R*^*i*^(*w*) satisfies 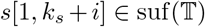, hence 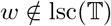. In words, 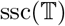 contains every taboo suffix *w* such that, for some *i* and every *i*-suffix *r*, string *wr* is the suffix of a taboo.

### Example 7

*If Σ*_1_ = {*A, C, G, T*} *and* 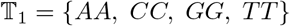, *then* 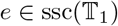, *for* 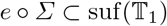.

## 8 The quotient graph

The quotient graph is a simple tool used to describe some properties related to the connectivity of graphs (see [3]).

### Definition 9

*Take a graph G* = (*V, E*) *and a partition of vertex set V, namely* 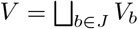 *for some index set J*.

*Then the* ***quotient graph of*** *G, denoted as Q*(*G*) = (*J, E*_*J*_), *is the graph whose vertices are J and such that* {*b*_1_, *b*_2_} ∈ *E*_*J*_ *iff an edge connects a vertex in* 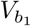 *with a vertex in* 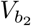.

See Figure 2 for a graphical example of a quotient graph. Our strategy to prove connectivity will be centered at the usage of following proposition, whose proof is simple enough to be omitted.

**Fig. 2.**
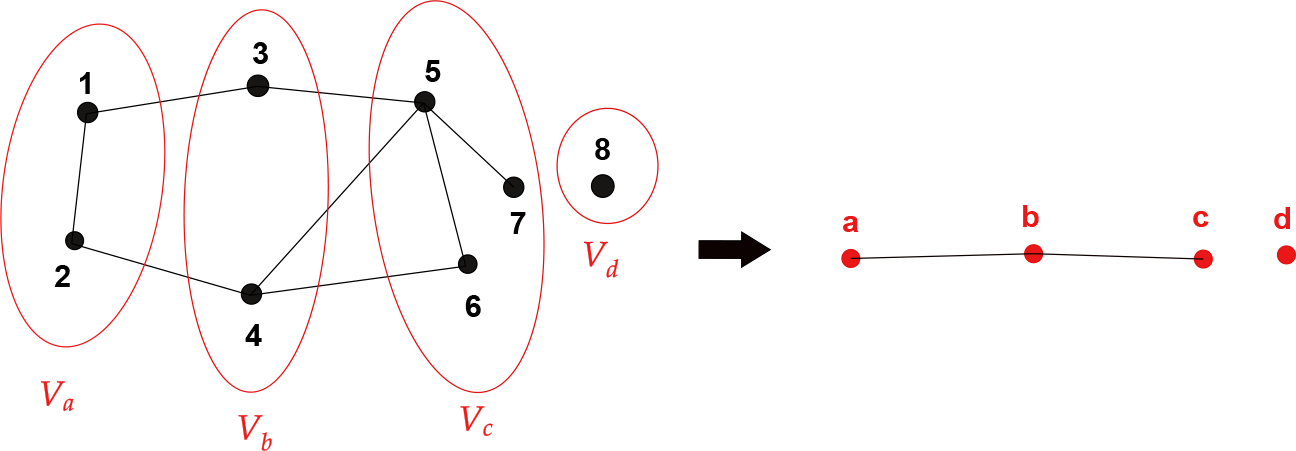
Example of a quotient graph. Given graph *G* = (*V, E*) on the left hand side, where *V* = {1, 2, 3, 4, 5, 6, 7, 8}, we consider partition 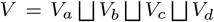, which induces the quotient graph *Q*(*G*) on the right hand side.

### Proposition 16

*Given graph G* = (*V, E*), *consider a partition* 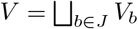.

*If every induced subgraph G*(*V_b_*) *is connected and the quotient graph Q*(*G*) *is connected, then G is also connected*.

The following is a similar result.

### Proposition 17

*Given graph G* = (*V, E*), *the following are equivalent:*

– *G is connected*.
– *For every partition of V, the quotient graph Q*(*G*) *is connected*.

Both Prop. 16 and 17 are basic, although productive results. The following proposition will highlight the strong interaction between the quotient graph and the synchronization of taboo-free strings.

### Proposition 18

*Given taboo-set* 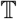, *j* ∈ ℕ_0_ *and n* ∈ ℕ_0_, *consider graph* 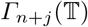,*a subset* 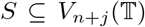 *and a partition* 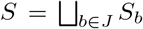. *Assume that* 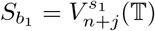 *and* 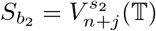 *for a pair of different b*_1_, *b*_2_ ∈ *J and different* 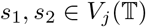 *consider the quotient graph* 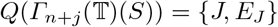.

*Then* {*b*_1_, *b*_2_} ∈ *E_J_ iff s*_1_ *and s*_2_ *are left n-synchronized and d*(*s*_1_, *s*_2_) = 1.

*Proof* By definition, *b*_1_ and *b*_2_ are adjacent in 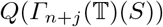 iff in graph 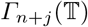 an edge connects a vertex in 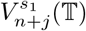 with a vertex in 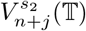. Since *d*(*s*_1_, *s*_2_) ≥ 1, this edge exists iff *d*(*s*_1_, *s*_2_) = 1 and there exists 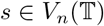 such that 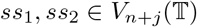. The last condition is the definition of *s*_1_ and *s*_2_ being left *n*-synchronized.

Prop. 18 combines very well with Prop. 7, where we proved that, **if** 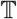 **is left proper** and strings *s, r* are left (*M* − 1)-synchronized, then *s* and *r* are left *k*-synchronized for any *k* ∈ ℕ_0_. Moreover, if *s, r* are left *k*^*^-synchronized, then they are left *k*-synchronized for any *k* < *k*^*^.

We now take 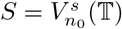 for some taboo-free string *s* and consider partition 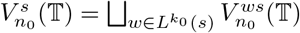 for some *k*_0_ ∈ ℕ with *n*_0_ ≥ *j* + *k*_0_. Then for two different 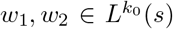, we set *s*_1_ := *w*_1_*s* and *s*_2_ := *w*_2_*s*. Prop. 18 implies that *w*_1_ and *w*_2_ are adjacent in 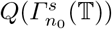 iff *s*_1_ and *s*_2_ are are left (*n*_0_ −*j*−*k*_0_)-synchronized and *d*(*w*_1_, *w*_2_) = 1. If *n*_0_ − *j* − *k*_0_ = *M* − 1, then we infer that *w*_1_ and *w*_2_ are left *k*-synchronized for arbitrary *k*, thus *w*_1_ and *w*_2_ are adjacent in 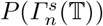 for arbitrary *n* ≥ *j* + *k*_0_. All in all, we have the following lemma.

### Lemma 19

*Given a left proper taboo-set* 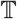, *j* ∈ ℕ_0_, 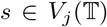 *and k* ∈ ℕ,*consider, for any n* ≥ *j*+*k*, *partition* 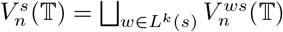 *and quotient graph* 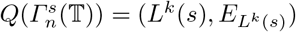. *Then it holds that*

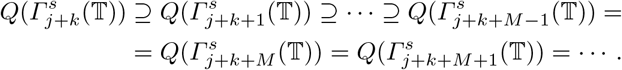

*If* 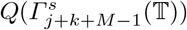 *is connected, then* 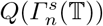) *is connected for n* ≥ *j* + *k*.

*Proof* We apply the mentioned *k*-synchronization properties and Prop. 18. If *n*_1_ > *n*_2_ ≥ *j* + *k*, then 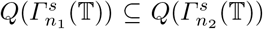. If *n*_1_ ≥ *j* + *k* + *M* − 1 and *n*_2_ ≥ *j* + *k* + *M* − 1, then 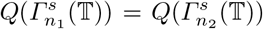. Regarding connectivity, given graphs *G*_1_ and *G*_2_ with the same vertex set *V*_1_ = *V*_2_ such that *G*_1_ ⊂ *G*_2_, if subgraph *G*_1_ is connected, then *G*_2_ is connected.

Figure 3 visualizes Lemma 19 for alphabet *Σ* = {*a, b, c*}, taboo-set 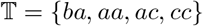 (which is left proper), suffix *s* = *b* and *k* = 1.

**Fig. 3.**
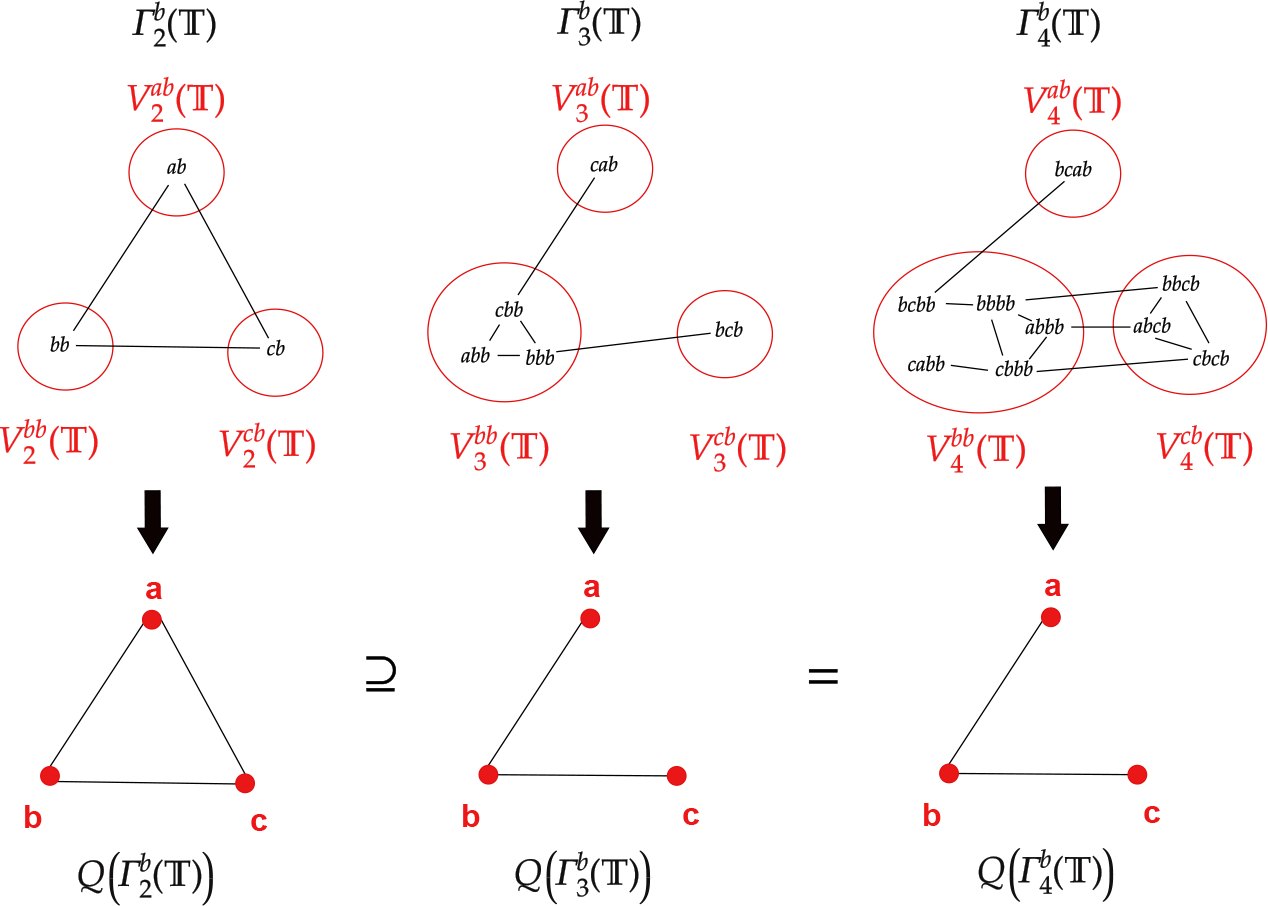
Visualization of Lemma 19 for *Σ* = {*a, b, c*}, 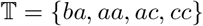, *s* = *b* and *k* = 1. It holds that *L*^1^(*b*) = {*a, b, c*}.

## 9 Connectivity of 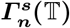 and 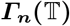

We are finally ready to study the connectivity of graphs 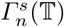 for |*s*| ≥ *M*.

### Lemma 20

*Given a left proper* 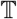, *for any* 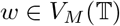 *consider the set* 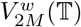 *and partition*

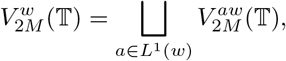

*inducing the quotient graph* 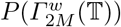. *Then the following are equivalent:*

a. *For every* 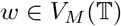, 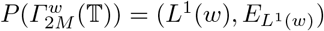 *is connected*.
b. *For every* 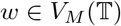 *and integer n* ≥ *M*, 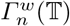 *is connected*.

*Proof* Prop. 17 states that, in a connected graph, every quotient graph is connected, thus *b*) implies *a*) by considering *n* = 2*M*. Let us prove by induction that *a*) implies *b*). For *n* = *M* and 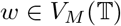, we have that

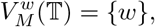

hence 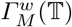 is connected. For the inductive step, assume that 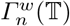 is connected for every 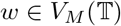 and up to an integer *n* ≥ *M*. We will prove that also every 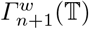 is connected. Consider

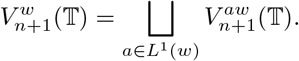

Let us write *w* separating the first *M* − 1 symbols from the last one, that is *w* = *rc* for *r* ∈ *Σ*^*M*−1^ and *c* ∈ *Σ*. Then for any *a* ∈ *L*^1^(*w*), 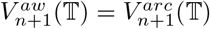. Since |*r*| = *M* −1, Prop. 9 implies 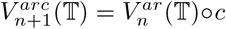, while the isomorphism of Prop. 13 yields

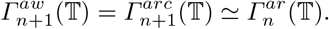

Thus every 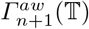 is connected, for the induction hypothesis implies that 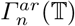 is connected since 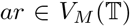. To prove that graph 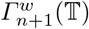 is connected, it remains to apply Prop. 16, so we need to prove that the quotient graph induced by partition 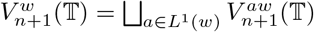, namely 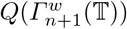, is connected.

We know that, given partition 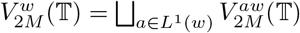, the quotient graph 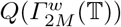 is connected. Applying Lemma 19 with *j* = *M*, *s* = *w* and *k* = 1, every 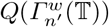 is connected for *n*′ ≥ *M*, as desired.

Lemma 20 is very interesting: We wanted to characterize the connectivity of graphs 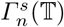 for 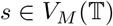 and *n* ≥ *M*. We have proved that it is enough to study a finite number of graphs, namely 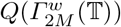 for 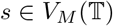, that is, 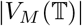 graphs. Let us state the connectivity results that follow from Lemma 20 and Theorem 15.

### Theorem 21

*Given a left proper* 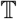 *the following are equivalent:*

a. *For any j ≥ M, any* 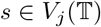 *and any integer n* ≥ *j*, 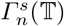 *is connected*.
b. *For any* 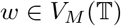 *and any integer n* ≥ *M*, 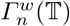 *is connected*.
c. *For any* 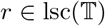, 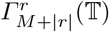 *is connected*.
d. *For any* 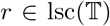, *partition* 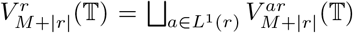, *induces a connected* 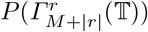.

*Proof* Implication *a*) ⇒ *b*) is obvious, while *b*) ⇒ *c*) is consequence of Theorem 15. Moreover, *c*) ⇒ *d*) follows from Prop. 17. To prove that *d*) ⇒ *a*), we split it into two parts, *d*) ⇒ *b*) and *b*) ⇒ *a*). Implication *b*) ⇒ *a*) follows from isomorphism 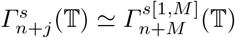 and 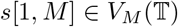.

It remains to prove *d*) ⇒ *b*). Corollary 11 implies *L*^1^(*w*) = *L*^1^(*w*[1*, k*_*w*_]). For any 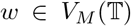 and *a, b* ∈ *L*^1^(*w*), we claim that *aw* and *bw* are left *k*-synchronized iff *aw*[1*, k*_*w*_] and *bw*[1*, k*_*w*_] are left *k*-synchronized.

Indeed, the implication ⇒) is obvious, so let us prove ⇐). Given a taboo-free string 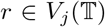 such that *raw*[1*, k*_*w*_] and *rbw*[1*, k*_*w*_] are taboo-free, we want to prove that also *raw* and *rbw* are taboo-free. But if that were not the case, it would be the consequence of either 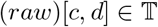 or 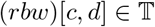 for some integers 1 ≤ *c* ≤ *j < j* + 1 + *k*_*w*_ ≤ *d* ≤ *j* + 1 + *M*. However, that contradicts the maximality of *k*_*w*_, yielding ⇐).

This claim and Prop. 18 imply that, if we write *r* = *w*[1, *k*_*w*_] for some 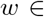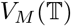, given partition 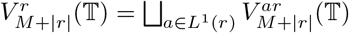, it holds that

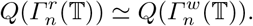

Theorem 15 implies that, for every 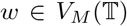, there exists 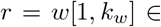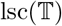. Applying Lemma 20, finally *b*) ⇒ *d*) follows.

It is worth noticing how simpler the connectivity problem has become. Initially, we were studying whether every 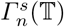 for 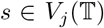 and *j ≥ M* is connected, obtaining in Lemma 20 that this is equivalent to the connectivity of graphs 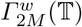 for 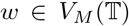, which are 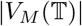 graphs. Now we see, using Theorem 21, that we only need to prove the connectivity of 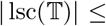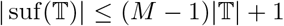 graphs, namely either 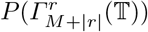 or 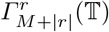 for 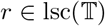. We give an example.

### Example 8

*Take Σ* = {*A, C, G, T*} *and* 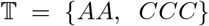, *which is left proper. Since M* = 3 *and* 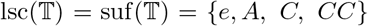, *the connectivity of graphs*

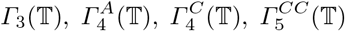

*implies that any* 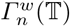 *with* 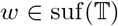 *is connected. Proposition 14 implies that, for any taboo-free string s and n* ≥ |*s*|, 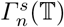 *is connected*.

Theorem 21 characterizes the connectivity of every 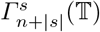 for |*s*| ≥ *M*. We know from Theorem 15 that there exists 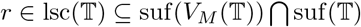 such that 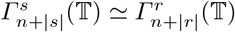. Since 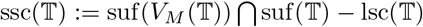, to complete our characterization of the connectivity of every Hamming graph with taboos, it remains to consider the connectivity of graph 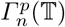 for 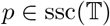. The techniques employed so far are flexible and give the following.

### Proposition 22

*Given a left proper* 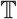 *and* 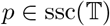, *assume that, for every* 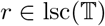, 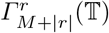 *is connected. Given k* ∈ ℕ, *if partition*

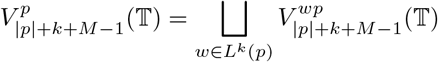

*satisfies that* 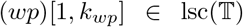 *for each w* ∈ *L*^*k*^(*p*)*, and moreover* 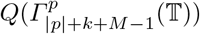 *is connected, then* 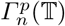 *is connected for n* ≥ |*p*| + *k*.

*Proof* For *n* ≥ |*p*| + *k*, given partition

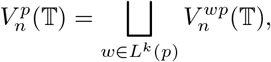

subgraphs 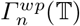 are connected due to 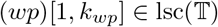. Moreover, since 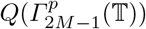 is connected, Lemma 19 with *s* = *p* implies that 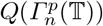 is connected for *n* ≥ |*p*| + *k*. The result follows applying Prop. 16.

In Prop. 22, one can always take *k* = *M* − |*p*| and just check if 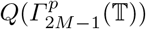 or 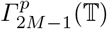 is connected. However, when 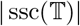 has just a few elements, it is easier to try a smaller *k*.

In general, if 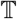 is not very restrictive, proving connectivity becomes easier since most of strings are left *k*-synchronized. In Prop. 23 only previous results are used, while in Prop. 24 we study this case more exhaustively in a self-contained manner. Note that, when taboo-set 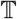 is minimal, the assumptions of Prop. 24 are much easier to check.

### Proposition 23

*Given a left proper* 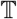 *such that every pair of strings* 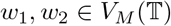 *with d*(*w*_1_*, w*_2_) = 1 *is left* 1*-synchronized, it holds that:*

a. *For any* 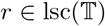 *and n* ∈ ℕ_0_, 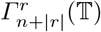 *is connected*.
b. *For any* 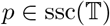 *with connected* 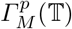, 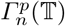 *is connected for n* ≥ *M.*

*Proof* Prop. 8 implies that every pair 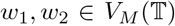 is left *k*-synchronized for any *k* ∈ ℕ. Any quotient graph 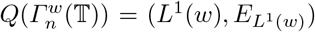 induced by partition 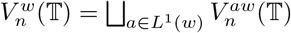 is completely connected, hence Theorem 21 implies *a*). Similarly with partition 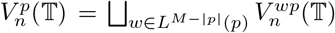, since 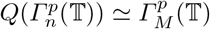 for *n* ≥ *M*, Prop. 22 imlies *b*).

### Example 9

*For Σ* = {*A, C, G, T*} *and* 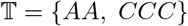, *the strings Tw*_1_ *and Tw*_2_ *are taboo-free for* 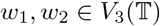, *hence they are left* 1*-synchronized. Since* 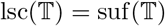, *for any taboo-free string s and n* ≥ |*s*|, 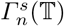 *is connected.*

### Proposition 24

*Given taboo-set* 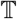 *and set* 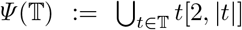, *if every pair of taboo-free strings* 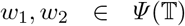 *with* |*w*_1_| ≥ |*w*_2_| *and d*(*w*_1_[1, |*w*_2_|], *w*_2_) ≤ 1 *is left 1-synchronized, then it holds that:*

a. *Every taboo-free string is* 1*-suffixable. In particular*, 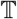 *is left proper.*
b. *Every two taboo-free strings s*_1_*, s*_2_ *with d*(*s*_1_*, s*_2_) = 1 *are left* 1*-synchronized.*
c. *Graph* 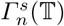 *is connected for every taboo-free string s and n* ≥ |*s*|.

*Proof*

a. Consider any taboo-free string *s*. Assume that, for each *a ∈ Σ*, *as* is not taboo-free, that is, that for some integer *c*_*a*_ ≤ 2, 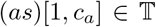. WLOG assume 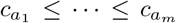 and consider 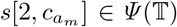, which is not left 1-synchronized with itself, what contradicts the assumptions of the statement, hence *s* is 1-suffixable. Taking 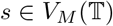 we see that 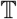 is left proper.
b. Given taboo-free strings *s*_1_*, s*_2_ such that *d*(*s*_1_*, s*_2_) = 1, assume that they are not 1-synchronized. Then for every *a* ∈ *Σ*, euither 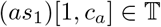 or 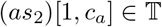 for some *c*_*a*_ ≥ 2. Denote by 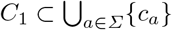 those *c*_*a*_ such that 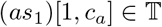, and analogously with *C*_2_. If *C*_1_ were empty, then *s*_2_ would not be 1-suffixable, contradicting *a*). Thus, both *C*_1_ and *C*_2_ must be nonempty. Consider *d*_1_ ≔ max{*c* : *c* ∈ *C*_1_} and *d*_2_ ≔ max{*c* : *c* ∈ *C*_2_}. It holds that 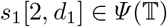 and 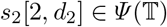. Moreover, we have that the pair *s*_1_[2*, d*_1_], *s*_2_[2*, d*_2_] is not left 1-synchronized. Since *d*(*s*_1_*, s*_2_) = 1, that contradicts the assumptions of the statement, hence *s*_1_ and *s*_2_ must be left 1-synchronized, as desired.
c. Clearly 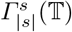 is connected, so let us proceed by induction. Assume 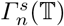 is connected for a fixed *n* ≥ |*s*| and consider 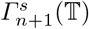. Since 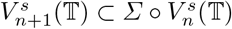, if 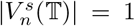, then 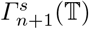 is connected. Otherwise we take different 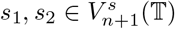; we will prove that they are connected. We know that 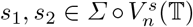, hence let us write *s*_1_ = *c*_1_*w*_1_ and *s*_2_ = *c*_2_*w*_2_ for *c_i_* ∈ *Σ* and 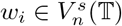. If *w*_1_ = *w*_2_, the result is obvious, so assume *w*_1_ ≠ *w*_2_.

By hypothesis, 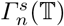 is connected, thus there exists a path of vertices of 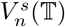, namely *y*_1_, · · ·, *y*_*D*_, such that *d*(*y*_*i*_, *y*_*i*+1_) = 1, *y*_1_ = *w*_1_ and *y*_*D*_ = *w*_2_. For every *j* ∈ [1, *D* − 1], the pair *y*_*j*_, *y*_*j*+1_ is left 1-synchronized, thus there exists *b*_*j*_ ∈ *Σ* such that *b*_*j*_*y*_*j*_ and *b*_*j*_*y*_*j*+1_ are taboo-free. Since *d*(*b*_*j*_*y*_*j*_, *b*_*j*_*y*_*j*+1_) = 1, *b*_*j*_*y*_*j*_ and *b*_*j*_*y*_*j*+1_ are adjacent in 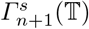. Moreover every pair of taboo-free strings contained in *Σ* ◦*y*_*i*_ is adjacent for *i* ∈ [1, *D*−1]. Since the relation “being connected” is transitive, vertices *s*_1_ ∈ *Σ* ◦ *y*_1_ and *s*_2_ ∈ *Σ* ◦ *y*_*D*_ are connected, as desired.

### Example 10

*If Σ* = {*A, C, G, T*} *and* 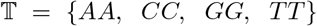, *then* 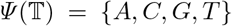. *Every pair of strings in* 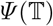 *is left* 1*-synchronized, hence for every taboo-free s and n* ≥ |*s*|, 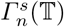 *is connected.*

Now we aim to find a lower bound for the number of taboos needed to disconnect one of the graphs 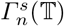. Let 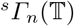 denote the subgraph induced in 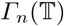 by all the taboo-free strings with **prefix** *s*. Then the following Corollary of Prop. 24 holds.

### Corollary 25

*Consider an alphabet Σ with m* ≔ |*Σ*| *and a taboo-set* 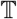. *The following holds:*

a. *If* 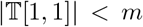, *then for any taboo-free string s and n* ≥ |*s*|, 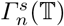 *is connected*.
b. *If* 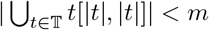, *then for any taboo-free string s and n ≥ s*, 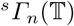 *is connected*.
c. *If* 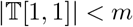 *or* 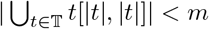, *then* 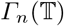 *is connected for n* ≥ 0.

*Proof*

a. Assume that taboo-free strings *s*_1_, *s*_2_ satisfy 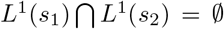. That is, for each *a ∈ Σ*, either *as*_1_ or *as*_2_ has a taboo as prefix, contradicting 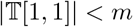. Therefore we can apply Prop. 24, implying *a*).
b. It follows from statement *a*), reversing the order of the symbols composing the strings.
c. Since 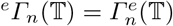, it follows from *a*) and *b*).

The upper bound 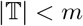, which guarantees connectivity for every 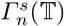, is the best one achievable without making additional assumptions. This is consequence of Corollary 25 and the following examples.

### Example 11

*If Σ* = {0, 1} *and* 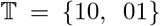, *then* 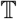 *is left proper and* 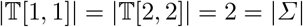. For 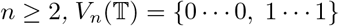, *what makes* 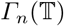 *disconnected. The trivial graphs* 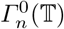 *and* 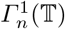 *are both connected.*

### Example 12

*For m* ≥ 3, *Σ* = {*a*_1_, · · ·, *a_m_*} *and the left proper taboo-set*

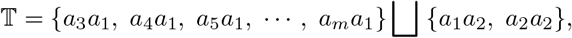

*we claim that* 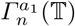 *is disconnected for n* ≥ 3. Indeed,

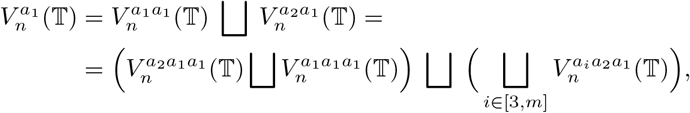

*so take* 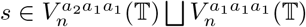 *and* 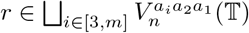. *It holds that d*(*s, r*) ≥ 2, *hence we found two disconnected components in graph* 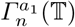. *This is coherent with* 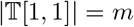. *Since* 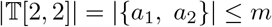, *Prop. 25 implies that* 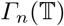 *is connected for any n* ∈ ℕ.

*To generalize this example, for i* ∈ ℕ_0_, *denote by* 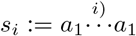 *the concatenation of i a*_1_*’s. The taboo-set*

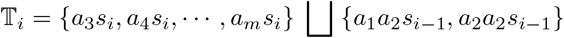

*satisfies that graph* 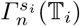 *is disconnected for n* ≥ *i* + 2*. It holds that* 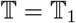, *where s*_0_ = *e and s*_1_ = *a*_1_.

## 10 Connectivity of the examples of Section 2

A permutation of the symbols of alphabet *Σ* does not alter any of the results that we proved along this work. Moreover, by reversing the order of the symbols, any statement regarding e.g. left-properness and suffixes has an analogous one in which right-properness and suffixes are involved. On the other hand, taboo-sets induced by restriction enzymes remain invariant when we interchange every recognition sequence by its reverse complement. Therefore, note that, for a bacterial taboo-set 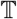, if we prove that every graph 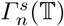 is connected, then also every graph 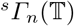 is connected.

### 10.1 Turneriella parva

Since 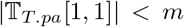, Corollary 25 implies that every graph 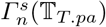 is connected. Therefore the evolution of the DNA sequences can potentially reach any other taboo-free DNA sequence, no matter which suffix was conserved along this process.

### 10.2 Helicobacter pylori

We want to apply Prop. 24. Take any 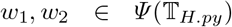 and assume that they are not left 1-synchronized. In particular WLOG we can assume that *T* ∉ *L*^1^(*w*_1_), implying *w*_1_ = *GCA*. If *C* ∉ *L*^1^(*w*_1_), then *w*_1_ ∈ {*CGG, CTC, ATG*}, which is absurd since it contradicts *w*_1_ = *GCA*. Therefore it must be *C* ∉ *L*^1^(*w*_2_), yielding *w*_2_ ∈ {*CGG, CTC, ATG*}. In any case, *d*(*w*_1_, *w*_2_) ≥ 2, so Prop. 24 can be applied, therefore every graph 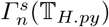 is connected. In particular, 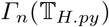 is connected.

### 10.3 An imaginary bacterium

Proposition 24 cannot be applied because *CCC* and *GCC* are not left 1-synchronized. Consistently, the graph 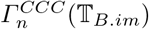 is disconnected for *n* ≥ 5. To prove this, take 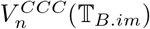, which satisfies

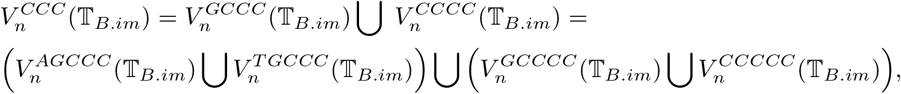

implying that, for any strings 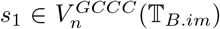 and 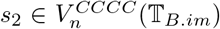, it holds that *d*(*s*_1_, *s*_2_) ≥ 2. Thus we found two disconnected components in 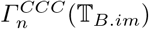, namely 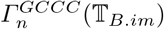 and 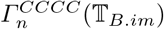.

## 11 Concluding remarks

Using the results proven in this work, it is possible to decide whether every Hamming graph 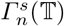 is connected. In most of biological cases, with just a small number of recognition sequences, it seems very likely that all these graphs are connected. This can be checked for most of instances using Prop. 24. In this sense, our work supports the assumption tacitly made when studying DNA-sequence evolution – namely, that every DNA sequence can reach any other sequence by single-point substitutions.

On the other hand, our formal framework is a first and necessary step to study the effect of restriction enzymes in the DNA composition of bacteria and virus. Consider, for example, the phylogenetic studies in [1], where our *H. pylori* taboo-set 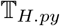 was taken from. Is the inferred evolutionary time between two *H. pylori* populations affected by 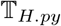? Has their *GC* content varied due to the taboos of restriction enzymes? To give an answer, the random mutation of DNA can be modelled in a Monte Carlo framework, although the connectivity of graphs 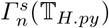 is of fundamental importance.

More in general, recent progress has been made in the usage of restriction enzymes for the treatment of viral infections ([23]). Since one or just a few SNPs can significantly alter the symptoms or even the mortality associated to a virus infection (cfr. [4], [25], [15]), the techniques employed here could help to identify which adjacent SNPs hinder the harmful ones by inducing the cleave of the restriction enzymes. Although such strategies are currently unexplored, this work could be a first theoretical guide to a successful treatment.

## 13 Authors’ contributions

Cassius Manuel wrote this manuscript with the guidance of Arndt von Haeseler. All authors read and approved the manuscript.

## 14 Acknowledgements

We would like to thank Michael Charleston (University of Tasmania) for his constructive criticism. In fact the idea of studying taboo-free sequences originated from discussions with Mike while Arndt received a visiting fellowship at the University of Tasmania. This work was supported by the Austrian Science Fund (FWF, grant number I-1824-B22) to Arnd von Haeseler.

